# The Cancer Epitope Trees of 23 Early Cervical Cancers in Chinese Women

**DOI:** 10.1101/043927

**Authors:** Xia Li, Hailiang Huang, Yanfang Guang, Yuhua Gong, Chen-Yi He, Xin Yi, Ming Qi, Zhi-Ying Chen

## Abstract

Emerging evidences suggest the heterogeneity of cancers limits the efficacy of immunotherapy. To search for optimal therapeutic targets, we used whole-exome sequencing data from 23 early cervical tumors from Chinese women to investigate the hierarchical structure of the somatic mutations and the predicted neo-epitopes based on their strong binding with major histocompatibility complex class I molecules. We found each tumor carried 117 mutations and 61 neo-epitopes in average and displayed a unique phylogenic tree and “cancer neo-epitope tree” comprising different compositions of mutations or neo-epitopes. Conceivably, the neo-epitopes at the top of the tree shared by all cancer cells are the optimal therapeutic targets that might lead to a cure. Human papillomavirus can be used as therapeutic target in only a proportion of cases where the integrated genome exits without active infection. Therefore, the “cancer neo-epitope tree” will serve as an important source to determine of the optimal immunotherapeutic target.

## Introduction

Immunotherapy is emerging as the most promising type of cancer treatment, as evidenced by recent clinical trials in which durable remission, and even cure, have been demonstrated in some patients. However, its success is limited because only a small proportion of patients respond to the therapies, whereas most remain resistant and are either unresponsive or responsive only transiently^1, 2^. A growing body of evidence suggests that cancer heterogeneity is the bottleneck that limits the efficacy of immunotherapy. Although derived from a single initiated cell, almost all cancers comprise multiple subclones and evolve constantly, driven by mechanisms such as genome instability and Darwinian selection. Some recent cancer genome studies have revealed that subclones of a cancer comprise different compositions of genomic alterations^3, 4^. It is conceivable that some of these alterations have become the determinants of whether the subclone responds to or resists current immunotherapies, including immune checkpoint inhibitors, tumor infiltration lymphocytes, chimeric antigen receptor modified T cells, and bispecific antibodies. Each of these immunotherapy strategies targets only one or a few subpopulations and allows the others, especially the metastatic ones, to continue to thrive^5^. It is also conceivable that these genetic variabilities have become the bottleneck that limits clinical efficacy; many patients either have no response or have an incomplete response in which the cancer shrinks or even disappears but eventually recurs despite continuing treatment.

It has become apparent that the development of technology to target all of a tumor’s subclones is necessary to remove the bottleneck and bring cancer immunotherapy to a new level. We recently proposed the construction of a “cancer epitope tree” to achieve this goal^6^ because, although the cancer genome is highly variable both spatially and temporally, the technology is available to determine the subclonal hierarchical structure, the so-called phylogenetic tree, to outline the temporal relationship among the genomic alterations from various subclones^7^, even at the single-cell level^8^. In combination with the technique to predict the neo-epitopes created by these genomic alterations, a “cancer neo-epitope tree” can be constructed to guide the systematic search for the optimal therapeutic targets located at the “trunk” or “major branch” that possess the potential to mediate T-cell killing of all or most of the cancer cells^6^. With few exceptions^9^, most cancer phylogenetic trees were constructed with driver mutations at a cohort level. Passenger mutations outnumber driver mutations by up to 2,000 times^3, 10, 11^; many of them are target candidates and could play an important role in causing cancer cell death by cytotoxic T lymphocytes and antibody-dependent cell-mediated cytotoxicity. Therefore, for purposes of immunotherapy, it is important to include neo-mutated epitopes derived from both driver and passenger mutations. Furthermore, an accumulating body of evidence suggests that each cancer is unique in its composition of genetic alterations and hierarchical structure of subclones^7, 9^. Therefore, a “cancer epitope-tree” of an individual cancer will be more useful than a cohort level for determination of immunotherapeutic targets.

In this article, we report the results from the first attempt to construct the individual “cancer epitope tree” of 23 early cervical tumors in Chinese women. Cervical cancer is the most lethal cancer in women worldwide, with an estimated 528,000 new cases and 266,000 deaths in 2012^12^. Persistent infection with high-risk human papillomavirus (HPV) subtypes, such as HPV 16 and 18, has been found in a majority of patients with cervical cancer. Although multiple studies have characterized the mutation landscape of cervical cancer^13, 14^, the molecular events responsible for malignant transformation remain elusive. Because the 23 cervical tumors in this study were at very early stages (17 stage I and 6 stage II, according to the International Federation of Gynecology and Obstetrics [FIGO] staging system), they allowed us to study the early events in cervical carcinogenesis while defining the optimal immunotherapeutic targets. We found that the samples from 73.9% of our patients with early-stage cervical cancer carried the integrated HPV genome. The ubiquitin proteolysis and extracellular matrix (ECM) receptor pathways were significantly altered; 69.6% of cancers carried alterations in these two pathways. We propose that alterations in genes *FBXW7* and *PIK3CA* are more likely to serve as the early genomic mutations that cause the progression of HPV-induced precancerous cells toward invasive malignancy. Furthermore, we used the identified somatic mutations to predict the neo-epitope on the basis of their affinity with major histocompatibility complex class I molecules (MHC-I). Using this information, we constructed the phylogenetic tree and the “cancer epitope tree” for individual tumors. We found that the mutations of individual tumors displayed a unique path of evolution, highlighting its importance in the search for therapeutic targets. HPV proteins might serve as immunotherapy targets in tumors that carry the integrated virus genome without active HPV infection. However, for tumors that do not express these proteins, our approach will suggest desirable therapeutic target candidates. The results of this study expanded our understanding of the early stages of cervical carcinogenesis and, more importantly, offered a useful systematic strategy with which to search for the optimal immunotherapeutic targets, which has important implications for cancer diagnosis, prevention, and therapy.

## Results

### General data

Our study included 23 patients from the Southwest Hospital of Chongqing Autonomous Municipality in China who had received a diagnosis of early-stage cervical cancer (17 stage I and 6 stage II, according to the FIGO staging system; Table 1). We performed whole-exome sequencing of 242,232 exons, with a length of 63.8 megabases, at an average coverage of 181X. Peripheral blood samples from most patients were used as germline controls, with the exception of patients S21, S22, and S23, for whom adjacent tissues were used instead. We detected HPV sequences in all but two tumor samples and found genomic integration only in 17 exome-captured sequencing datasets (Supplementary Table 1). We used MuTect and Indelocator^15, 16^ to call each case’s somatic mutations by filtering out germline events from the corresponding normal sample. We also filtered out variants in the 1000 Genome Project (Phase 3)^17^, the NHLBI GO Exome Sequencing Project (version 2)^18^, and the Exome Aggregation Consortium (version 0.2)^19^ by applying a minor allele frequency threshold of 0.1 to all three databases. The final cleaned dataset includes 2,691 somatic mutations, including 730 synonymous substitutions, 1,934 nonsynonymous substitutions, 18 deletions, and 9 insertions across 23 sample pairs. A subset of 59 somatic mutations was selected for validation, and 57 variants (96.6%) were validated using mass spectrum or Sanger sequencing (Supplementary Table 2). The number of nonsynonymous mutations show no correlation with the patients’ age or clinical stage (Supplementary Fig. 1).

**Table 1.**
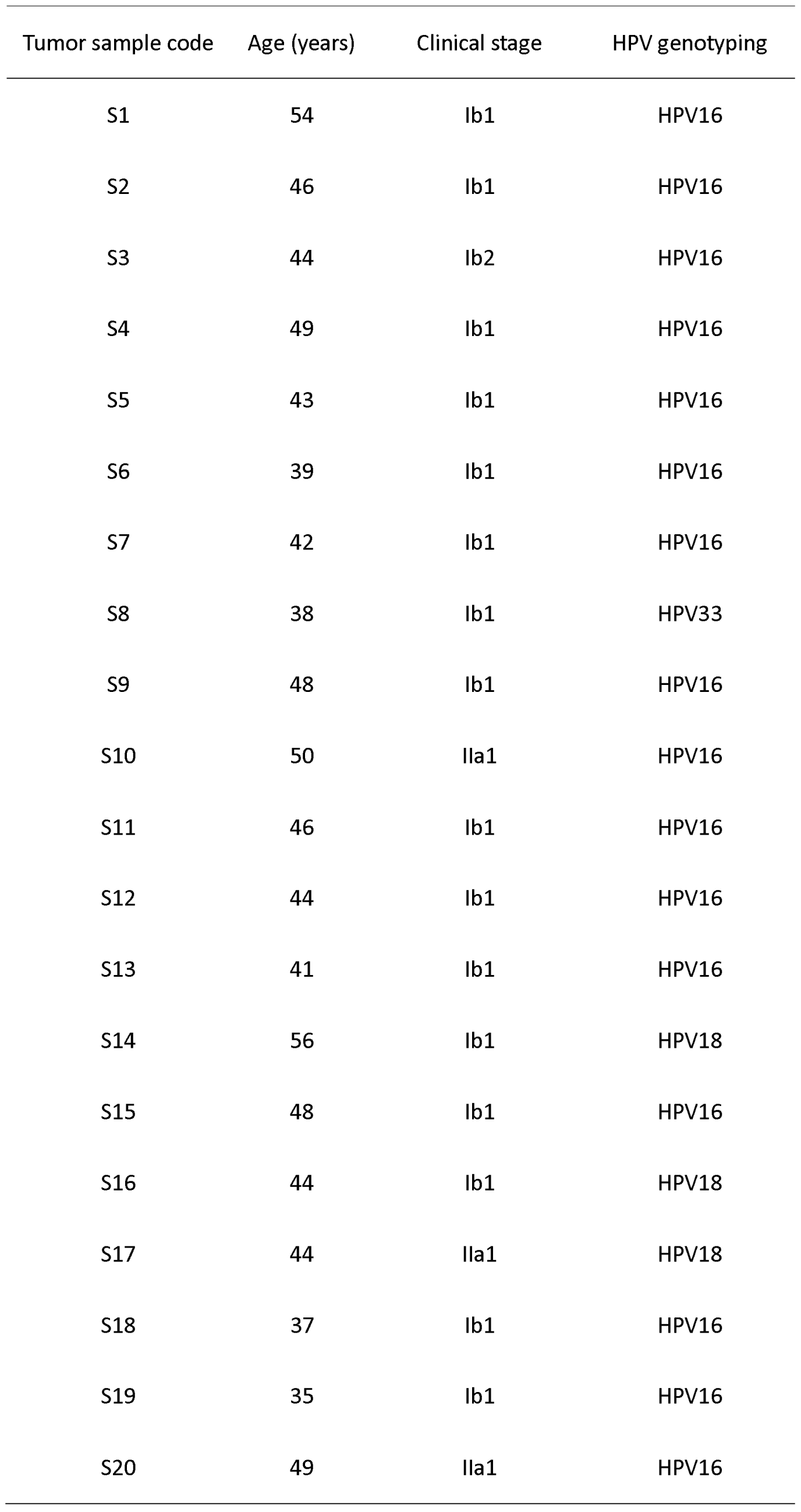
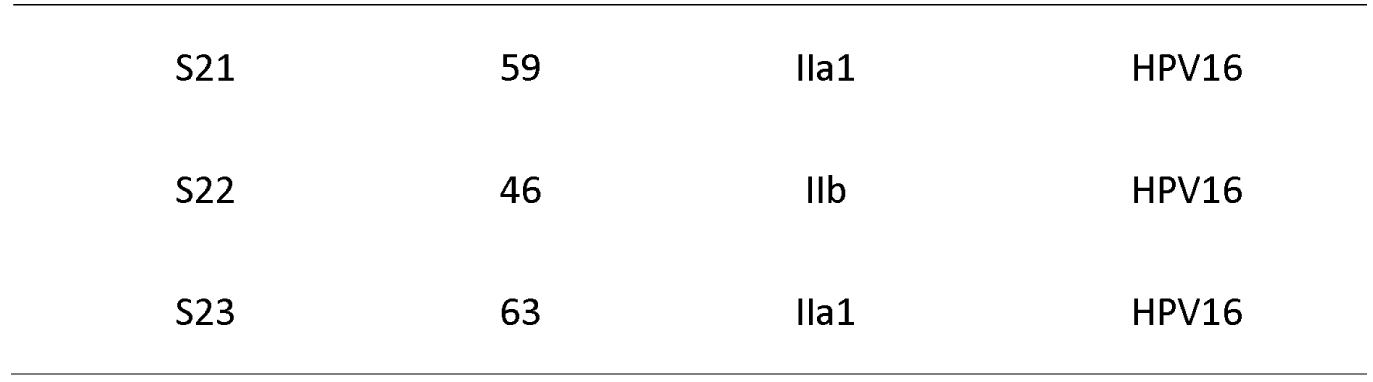
Clinical stage and HPV infection status of 23 patients with cervical cancer.

### Frequency of Mutations in Cervical Cancer

We first estimate the distribution of the somatic mutations and their nucleotide substitutions. C/T and G/A substitutions were the most frequent among the 23 patients (Fig. 1b), with mean frequencies of 21.8% and 21%, respectively. This observation, especially the C/T substitution pattern, agrees with findings from a previous study of 115 Norwegian and Mexican cervical cancer samples^13^. We found that *PIK3CA* (17.4%), *SYNE1* (17.4%), *FBXW7* (17.4%), and *MUC16* (21.7%) were among the most frequently mutated genes (Fig. 1a), which again agrees with the findings of previous studies: the Norwegian and Mexican study^13^ showed that *EP300* (16%), *FBXW7* (15%), and PIK3CA(14%) harbored recurrent mutations; a study^20^ in 80 cervical cancer samples from Boston showed that *PIK3CA* (31.3%), *KRAS* (8.8%), and *EGFR* (3.8%) had the highest mutation rates; and another study in 15 cervical cancer patients from Hong Kong revealed frequent alteration of *FAT1* (33.3%), *ARID1A* (33.3%), *ERBB2* (26.7%), and *PIK3CA* (53.3%)^21^. Despite the ethnic and geographic differences, alterations in *PIK3CA*, followed by *FBXW7*, were the most common mutations in the various cervical cancer studies.

**Fig 1.**
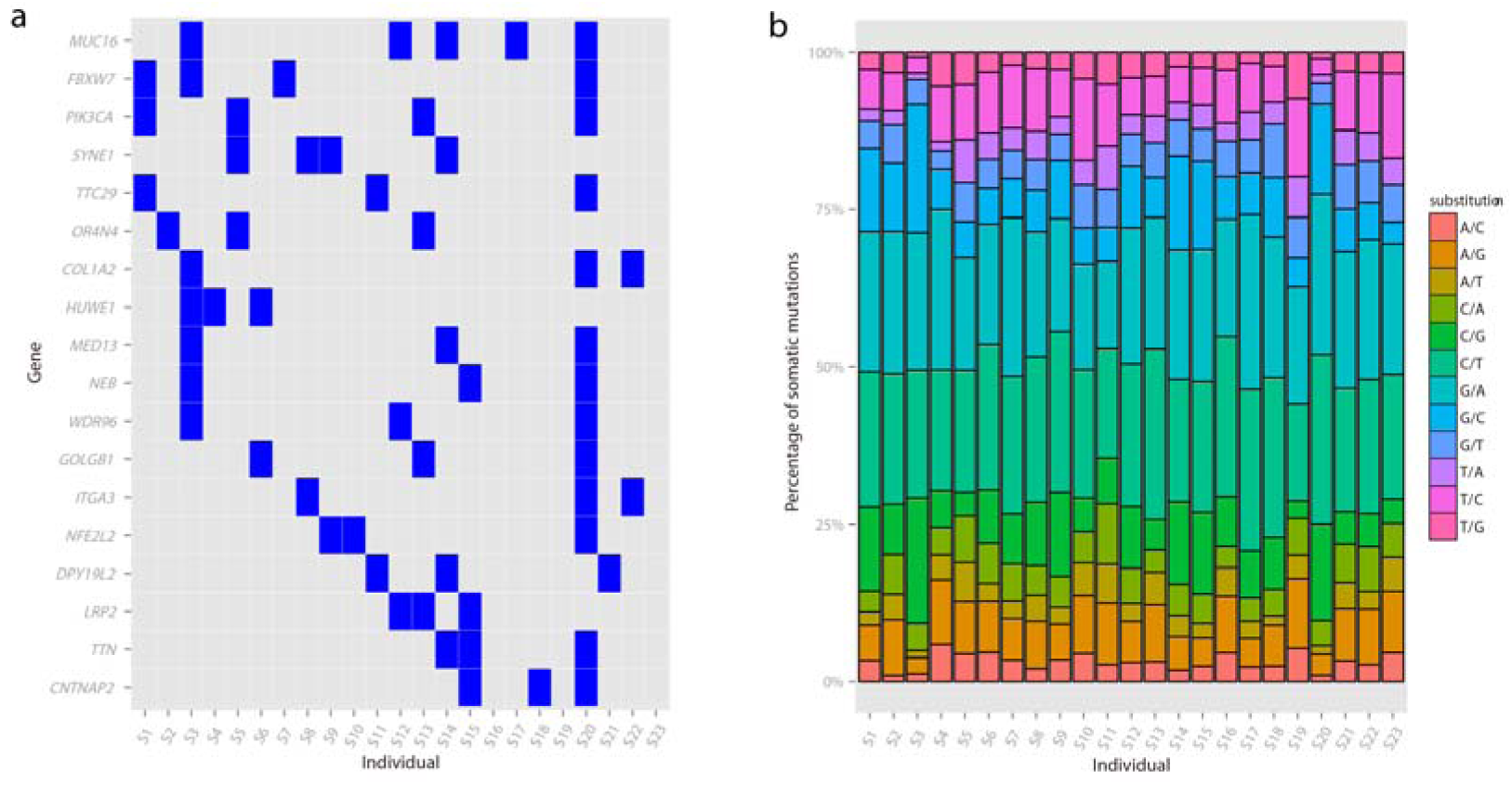
Distribution of mutated genes and base substitution patterns in the 23 patients with cervical cancer. (a) Distribution of mutated genes in at least three patients (mutation frequency > 13%). Each column represents one individual, and each row is a gene. (b) Distribution of base substitution patterns for all somatic mutations.

### Identification of neo-epitopes

The neo-antigen identification approach estimates the binding affinity between the mutated peptide and MHC-I and can be used to predict potential targets for immunotherapy^22, 23, 24, 25, 26^ Defining the immunogenic peptides that have strong binding affinity with MHC-I when mutated (see Methods), 1,405 of the 1,934 nonsynonymous substitutions showed immunogenicity (Fig. 2), which suggests that tumor progression generates antigens that may recruit immunologic cells to attack the tumor cells.

**Fig 2.**
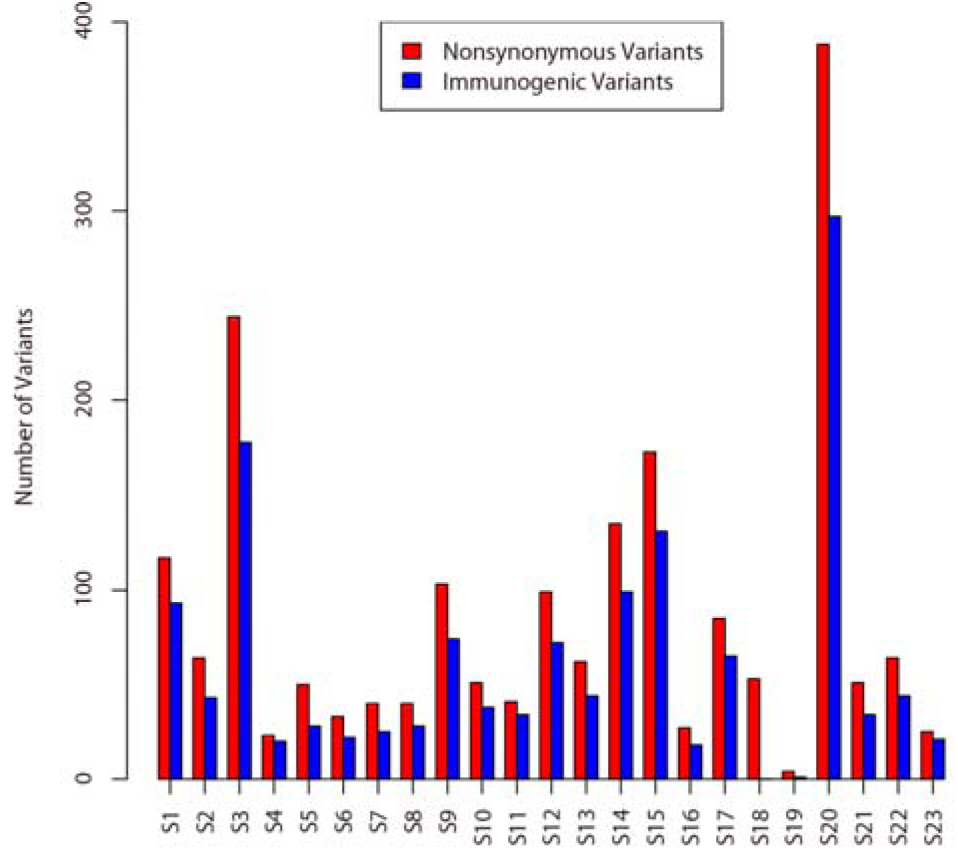
Number of immunogenic variants in 23 patients. The numbers of total nonsynonymous mutations (red) and immunogenic variants (blue) are shown for each patient. The variant showed immunogenicity only if the mutated peptide showed strong binding affinity with MHC-I (affinity < 50) and the normal peptide had no binding affinity (affinity > 500) at the same peptide position.

### Alteration in ubiquitin-mediated proteolysis and ECM receptor interaction pathways

To determine whether any biological functions were strongly enriched, we integrated all of the nonsynonymous mutations from the 23 patients and determined the pathways that were enriched. In permutation tests of 10,000 samples among the 177 mutated pathways, the ubiquitin-mediated proteolysis and ECM receptor interaction pathways were the most significantly altered, with false discovery rates of less than 0.1 (Table 2). All of the mutated genes involved in these two pathways are shown in Supplementary Fig. 2.

**Table 2.**
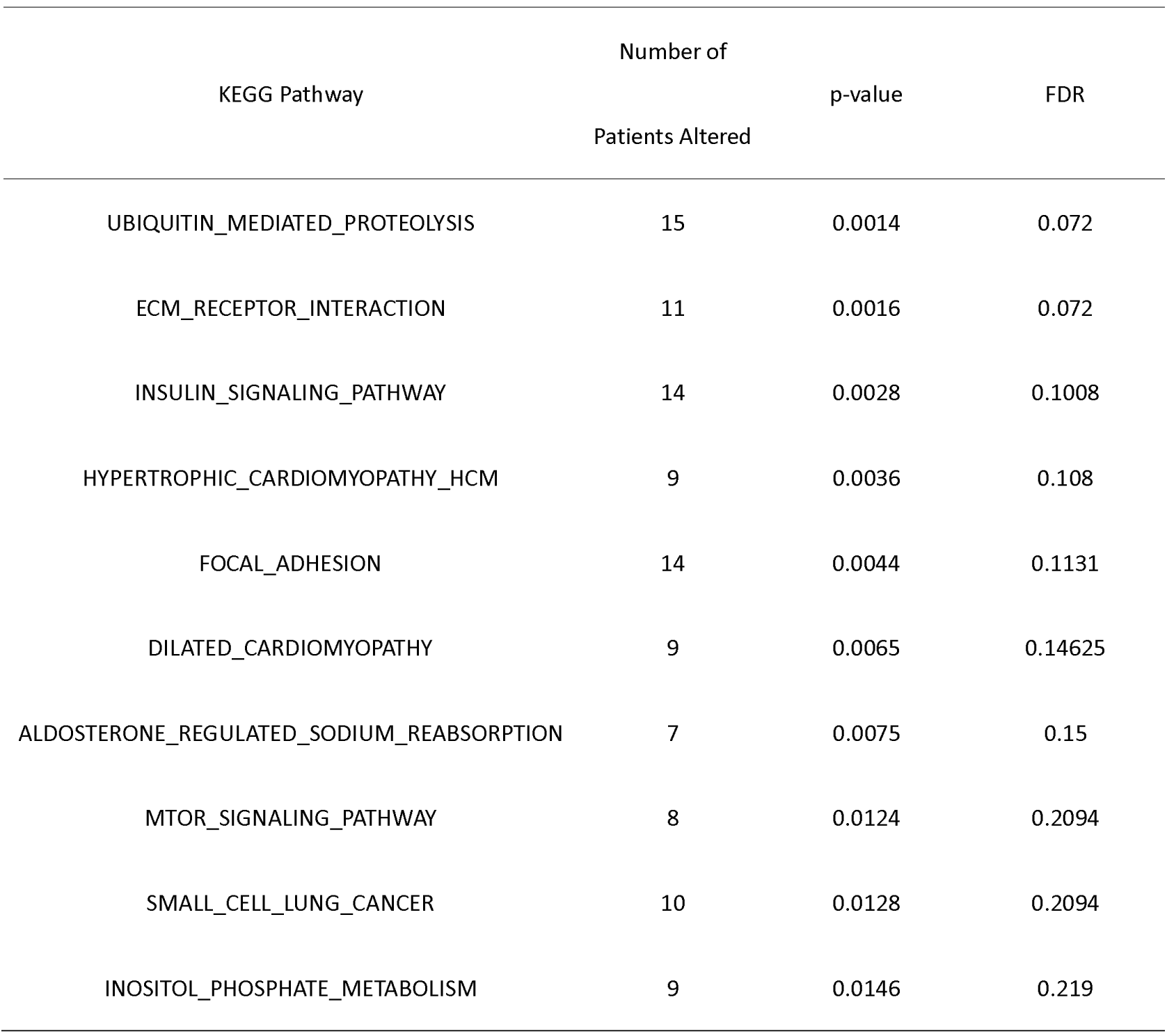
Top 10 altered pathways.

The ubiquitin-mediated proteolysis pathway mediates protein degradation via the ubiquitin conjugation and proteasome system. It is reported to be the most frequently altered pathway in clear cell renal cell carcinomas and a contributor to the tumorigenesis^27^. In our cervical cancer data, the mutated genes involved in this pathway included *FBXW7* (altered in 17%), *HUWE1* (altered in 13%), and *BIRC6* (altered in 9%). It has been suggested that mutations in *FBXW7* cause increased genetic instability because several prominent oncogenes (*Notch*, *c-Myc*, *JunB*, and *mTOR*) are its substrates^28, 29^. In cervical cancer, the ubiquitin-mediated proteolysis pathway can be best characterized by high-risk HPV-16 E6 binding activity to the tumor-suppressor protein p53 to induce ubiquitylation and proteasomal degradation^30, 31, 32^, and the abrogation of p53 allows the accumulation of genetic mutations that would normally have been repaired. The HPV-18 E7 oncoprotein also targets the tumor suppressor *Rb* proteins for proteasomal degradation via the ubiquitin-dependent pathway^33, 34^. Although the mTOR signaling pathway is not the most significant, it is among the top 10 altered pathways (with *PIK3CA* altered in 17% of patients). Thus, the alterations in genes involved in the ubiquitin-mediated proteolysis pathway may trigger a cascade of reactions that lead to malignancy.

For the ECM receptor interaction pathway, genetic alterations mainly occurred in *COL1A2* (altered in 13%) and *ITGA3* (altered in 13%), which may disrupt the signaling transfer function during interactions with extracellular proteins and therefore lead to malfunction in cellular activities such as adhesion, migration, differentiation, proliferation, and apoptosis. This pathway showed a certain overlap with the focal adhesion pathway involving mutations in *PIK3CA* (altered in 17%), *COL1A2* (13%), and *ITGA3* (13%). HPV-positive cells have been found to express high levels of focal adhesion kinase, which regulates the interaction between the signal transduction of ECM and integrins^35^. The virus oncoprotein HPV-16 E6 also binds to the ECM protein leading to cytoskeletal reorganization and formation of focal adhesions^36^. This interaction, in combination with de-regulation of focal adhesion kinase, promotes resistance to anoikis and allows the HPV-infected cells to proliferate in the absence of adherence to the ECM, i.e., anchorage-independent growth^37^. The altered genes in these pathways may allow cells to escape anoikis and play a role in transformation and tumor invasion.

### Evolutionary paths in individual patients

Tumors usually contain multiple genetically diverse clones or subclones that have constantly evolved from an earlier population through expansion and selection^3, 38, 39^. Outlining the evolutionary history of these mutations will aid in understanding the cancer development and guide design of therapy targets^40, 41^. We therefore constructed phylogenetic trees for each tumor using nonsynonymous substitutions (Fig. 3, Supplementary Table 3) and named the clones in chronical order as the ancestor, descendant, and later subclones. Consistent with their early stage of malignancy, the subclonal hierarchy structures of all tumors were simple. Five tumors (S4, S9, S11, S12, and S21) harbored only one ancestor subclone, and no descendants were observed. The evolutionary paths in the other tumors showed either linear (S3, S7, S8, S19, and S20) or branching (the remaining 13) patterns. Five of the 13 tumors with branching paths (S6, S13, S15, S16, S23) had two ancestor subclones (S6 and S15 derived one descendant subclone from one of the two ancestors), and the other eight carried only one ancestor subclone with multiple descendants or later subclones (Fig. 3).

**Fig 3.**
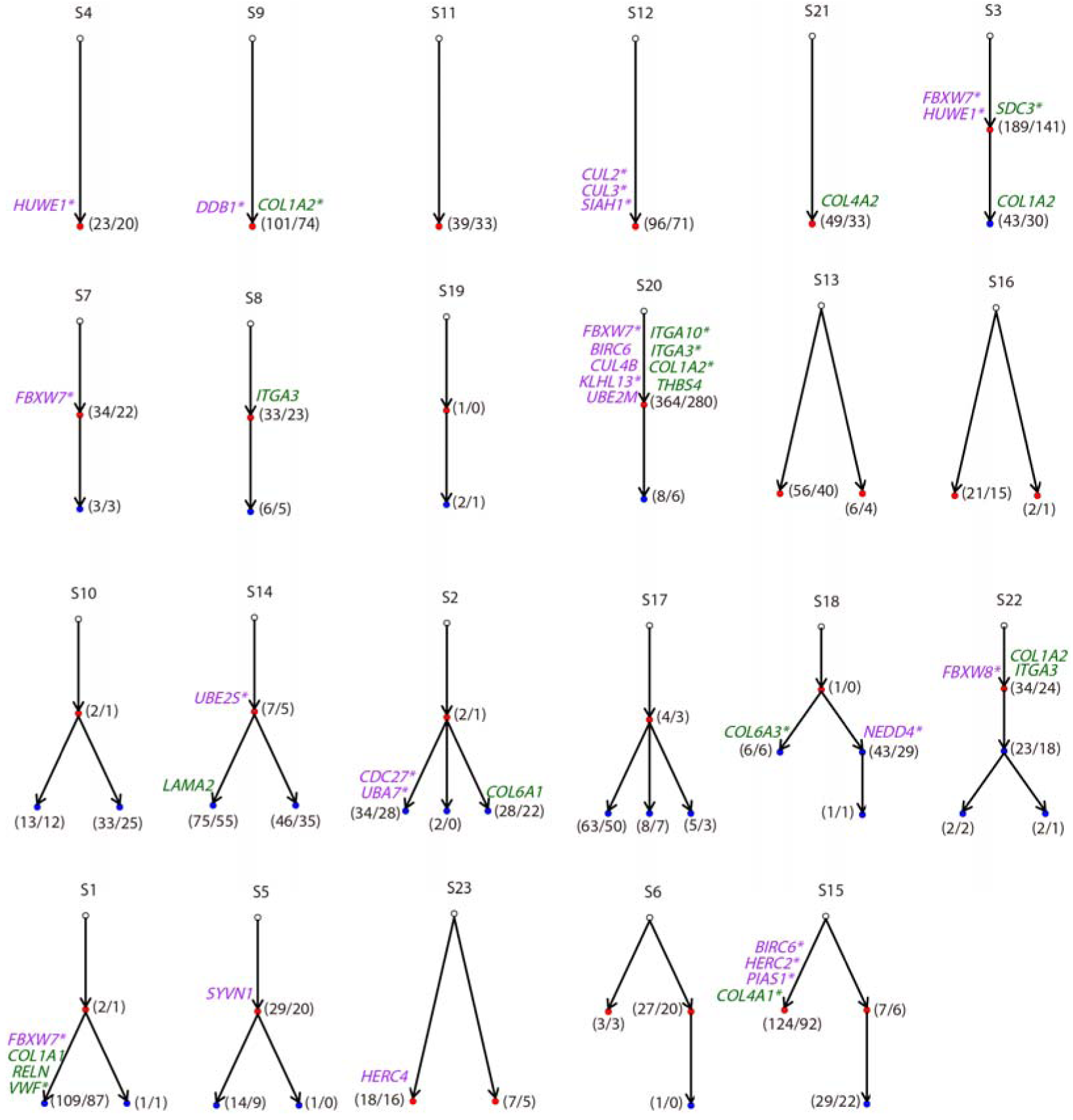
The clonal structures and phylogenic relationships for all 23 patients. In each patient, a phylogenic tree was constructed using somatic mutations. Each node represents one clone. Each clone harbors multiple mutations, and only the genes involved in the ubiquitin-mediated proteolysis (purple) and ECM receptor interaction (dark green) pathways are labeled on the corresponding node. Normal clones (non-pathogenic) are in open circles, early clones are shown in red, and later clones are shown in blue. Arrows point from the parent node to the child node (i.e., the descendant clone derived from the ancestor clone). Asterisks indicate that the gene harbors neo-epitopes. The numbers in parentheses indicate the number of total mutated genes (before the slash) and the number of genes that harbored neo-epitopes in the clone.

Each tumor displayed accumulation of different mutations and evolutionary paths over time, suggesting heterogeneity between patients. Thus, for therapeutic considerations, the individual phylogenetic tree should offer clues for the selection of therapy targets. The number of altered genes in each subclone and the number of genes that harbored neo-epitopes are shown in Fig. 3. Each tumor carried an average of 117 mutations and 67 antigenic targets. All tumors but S19, which only harbored three mutations, had subclones that harbored neo-epitopes, which makes immunotherapy a feasible approach. An individualized “cancer epitope-tree” could be constructed using neo-epitopes. Selection of targets in the ancestor subclones would inhibit the majority of the tumor cells because the descendants are derived from the ancestor subclone. We defined the ancestor subclones as the “trunk” and the descendant subclones as the “major branches” in the phylogenic tree. Among the many neo-epitopes in the trunk and major branches, one possibility to choose functional mutations or to scale down the mutation is to choose the genes involved in important pathways. We therefore list in Fig. 3 the 34 altered genes involved in the ubiquitin-mediated proteolysis and ECM receptor interaction pathways in these trees, together with the number of neo-epitopes.

### Alteration of *FBXW7* and *PIK3CA*

Mutations of both passenger and driver genes occur during a lesion’s transition from precancerous to malignant. Among the approximately 20,000 protein-coding genes in the human genome, only 138 genes were reported in a previous study as driver genes^42^, which play a significant role in tumorigenesis. We found that 24 of the 138 proposed driver genes were mutated in our 23 tumors (Fig. 4). In addition to *FBXW7* (17%) and *PIK3CA* (17%), which were the most frequent, *NFE2L2* (13%) and *CREBBP* (9%) were also frequently mutated. *NFE2L2* participates in protein processing and amino acid metabolism and was recently identified in a recent cervical cancer study^13^. Interestingly, in tumor S2, two driver genes, *GATA2* and *STK11*, were located in the ancestor subclone, and *STK11* also harbors neo-epitopes (Supplementary Table 3). It is possible that the two ancestral mutated driver genes granted a selective growth advantage to allow the cancer cells to derive more descendants in S2, as we observed. In the S2-specific “epitope tree,” *STK11*, which is part of the mTOR signaling pathway, may be a “trunk” target candidate.

**Fig 4.**
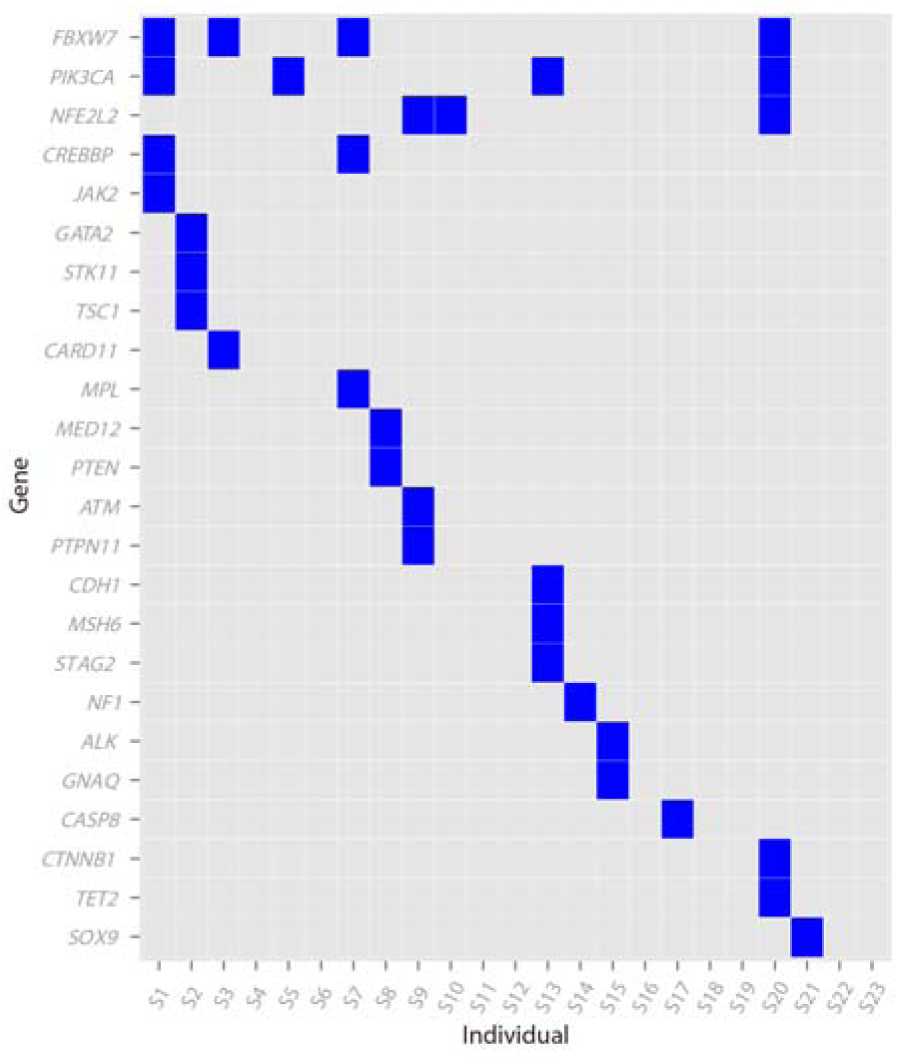
Distribution of the reported driver genes in our 23 patients. Driver genes were obtained from the literature^42^.

Overall, *FBXW7* and *PIK3CA* seem to play more important roles in these early-stage tumors. Both showed mutations in four patients. In the individual phylogenetic trees, both genes were located on the ancestor subclones in three tumors, which suggests they were likely early events during tumorigenesis. We propose that alterations in *FBXW7* and *PIK3CA* are likely the early changes that trigger the progression of the HPV-induced precancerous cells toward invasive malignancy. The other neo-epitopes involved in these pathways, located at the truck of the phylogenetic tree, might also be ideal candidates for immunotherapeutic targets.

## Discussion

Cervical cancer is among the few malignancies that allows convenient study from morphologic, cytological, and molecular events during the formation of precancerous lesions and their transition to invasive cancers. Consequently, timely diagnosis and treatment of early-stage cervical cancer is possible. The whole-exome sequencing data in this study were obtained from 23 patients with early-stage cervical cancer (FIGO stage I or II) to allow identification of early genomic events without the complication of late-stage genomic alterations. Infection of high-risk HPV is a prerequisite for cervical cancer, and integration of the viral genome occurs throughout the course of carcinogenesis. Twenty of the 23 patients had infection with high-risk HPV, and integrated HPV genomes were detected in 17 cases, albeit at relatively low sequences covered. A total of 2,691 genomic alterations, mostly single nucleotide substitutions, were identified, of which 1,405 were predicted to encode neo-epitopes on the basis of their strong binding affinity with MHC-I. Each cancer carried an average of 117 nonsynonymous somatic mutations and 61 neo-epitopes. To outline the phylogenetic relationship among these somatic mutations, we constructed the subclonal hierarchical structures of individual tumors and named the identified cancer cell populations in temporal order as ancestor, descendant, and late subclones. We found that five patients carried only the ancestor subclone, 16 carried an additional descendant, and only two had all three subclones. Furthermore, we found that 17% of the tumors had mutations in *PIK3CA* and *FBXW7* without mutation of typical driver genes, such as *KRAS*, *TP53*, and *EGFR*, as reported in other cervical cancer genome studies^13, 20^; these were not found in our study, which suggests that they may be the later-stage events.

It has been well documented that only HPV viral oncogenes E6 and E7 can induce precancerous lesions and that additional genetic alterations are required for malignant transformation^43^. Therefore, the evidence confirmed that most of the cervical cancers in this study were at a very early stage of malignancy. Therefore, the somatic mutations in our samples must have been capable of triggering the transition from benign to invasive lesions. Based on our analyses, we propose that mutation of *FBXW7* and *PIK3CA* and other members in these two pathways were among the earliest alterations that triggered malignant transformation. This hypothesis is consistent with earlier studies in a large number of cancer types in which mutation of *FBXW7* and *PIK3CA* was a frequent event, including cancers of the colon, brain, gastrointestinal system^44, 45^, cervix^21^, head and neck^46^, and breast^47^. This hypothesis is further supported by recent evidence that mutated *PIK3CA* initiates breast cancer by triggering multiple key events during the cancer initiation stage^48, 49^.

Our findings will be very useful in guiding cancer immunotherapy. A growing body of evidence suggests that cancer heterogeneity is the bottleneck that limits the efficacy of cancer immunotherapy^5, 50^. Most of the current immunotherapeutic technologies, including tumor infiltration lymphocytes, immune checkpoint inhibitors, chimeric antigen receptor-modified T cells, and bispecific antibodies, kill only one or a few subclones in a cancer and allow the others to continue to grow. In our study, we found that each of the 23 cervical tumors had a unique subclonal hierarchical structure that comprised a different composition of genetic alterations and neo-epitopes.

Therefore, the “cancer neo-epitope tree” of each tumor is critical to help determine the optimal targets at the trunk or major branch shared by all descendant cells that have the potential to lead to a cure. Another important observation is that a large number of passenger mutations encoded neo-epitopes that were potential target candidates. Many earlier studies also demonstrated that each cancer encodes a unique set of genetic alterations, but they focused on driver mutations and demonstrated the phylogenic tree at the cohort level without indicating their temporal relationship; thus, they are very useful in outlining cancer signal pathways, but not in determining the most suitable therapeutic target. Passenger mutations greatly outnumber driver mutations, so they may play an important role in cancer immunotherapy^51, 52^. Conceivably, the “cancer neo-epitope tree” strategy as established in this study will help to determine optimal therapeutic targets and result in a great increase in clinical efficacy or even cure, especially when a cocktail of targets is used to reduce the chances of escape due to sporadic loss of the targets.

In this study, we constructed the “cancer neo-epitope tree” using genomic data derived from a single DNA sample from each tumor. It should be noted that this approach is limited by many factors, such as the heterogeneous composition of the tumor’s cell population, the exome capture efficiency, the genomic sequencing and assembly technique, and the tumor cell collection method. Therefore, our technique is able to draw out a “cancer neo-epitope tree” that comprises only the major subpopulations. Even so, this is achievable only when the tumor is small and DNA is fully representative. For a large tumor, however, DNA from well-designed multiple samples^3, 11^ would be more appropriate.

In summary, our results show that each tumor carried a unique set of genetic alterations and associated epitopes and that the construction of individual “cancer epitope-trees,” together with the earliest genomic events, such as alterations in *FBXW7*, *PIK3CA*, and related members in the pathways, could assist in the systematic search for optimal targets at the trunk or major branches.

## Materials and Methods

### Sample collection and preparation

Twenty-three pairs of cervical cancer tumors and matched normal tissues were obtained from the Southwest Hospital of Chongqing Autonomous Municipality in China. The study protocol was approved by the Institutional Review Board of Southwest Hospital, and informed consent was obtained from each subject. Tumors and peripheral blood samples were collected from patients S1-S20, who each underwent surgical resection. For patients S21, S22, and S23, adjacent tissues were used as the control samples. The surgically resected tumors were snap frozen in liquid nitrogen and stored at ‐80. The blood samples were stored at ‐20. DNA was extracted from the frozen tissues and peripheral blood lymphocytes using commercial kits (TIANamp Blood DNA Kits and Genomic DNA Kits, Tiangen Biotech) and following the manufacturer’s instructions. HPV genotyping was performed using the polymerase chain reaction (PCR)-based mass spectrometry system^53^.

### Whole-exome sequencing

DNA from matched tumor and control samples were fragmented with an ultrasonicator UCD-200 (Diagenode). These fragments were purified and size selected with Ampure Beads (Beckman Coulter, Inc.) following three enzymatic steps (end repairing, the addition of an “A” base, and adapter ligation) according to Illumina’s instructions. NimbleGenEZ 64M human exome array probes (SeqCap EZ Human Exome Library v3.0) were used in hybridization. Each captured library was then pair-end sequenced in 100-bp lengths with an Illumina HiSeq 2000 following the manufacturer’s instructions.

### Read mapping and somatic mutation detection

Raw whole-exome sequencing reads were aligned to the reference human genome (hg19) using a BWA aligner (v 0.7.10)^54^ with default parameters. Alignments were sorted and converted into BAM format. Picard (v1.119) (http://picard.sourceforge.net/) was used to mark possible PCR duplicates in the BAM file, and the Genome Analysis Toolkit (v3.2.2)^55^ was used to improve alignment accuracy. Somatic point mutations were detected with MuTect (v1.1.4)^15^. Variants from the 1000 Genome Project (Phase 3)^17^, the NHLBI GO Exome Sequencing Project (version 2)^18^, which represented variants from more than 200,000 individuals, and the Exome Aggregation Consortium (version 0.2)^19^, which spanned variants from 60,706 unrelated individuals (with a minor allele frequency threshold of 0.1), were removed from the somatic mutations. Variants were annotated for effects on transcripts using the variant effector predictor tool^56^.

### Validation of somatic mutations

We validated a subset of recurrent mutations together with some randomly selected mutations by either mass spectrum or Sanger sequencing. Specific primers were designed for PCR amplification and base extension that covered the mutation sites. Genotyping assay and base calling procedures were performed on the MassArray platform of Sequenom by determining their genotypes in the tumors and matched samples. The PCR amplification products were sequenced with a 3730xl DNA Analyzer (Applied Biosystems). All sequences were analyzed with Sequencing Analysis Software Version 5.2 (Applied Biosystems).

### HPV genome alignment

The reads that could not mapped to the human reference genome were extracted and realigned to a database of multiple HPV reference genomes. HPV reference genomes were obtained from the Human Papilloma Virus Episteme (pave.niaid.nih.gov)^57^. With the paired-end-read information, we determined whether the HPV genome could integrate into the human genome by screening pairs of reads with one end mapped to the human genome and the other end mapped to the HPV genome.

### Pathway analysis

The KEGG pathways were obtained from the Molecular Signatures Database (MSigDB)^58^, and the gene set was downloaded from http://www.broadinstitute.org/gsea/downloads.jsp (accessed 19 Jun, 2015). The mutated genes in each tumor were compared with the KEGG pathway to determine whether the tumor had altered pathways.

For each pathway, we randomly sampled the same number of genes from all genes in the human genome without replacement. We then counted the number of tumors that harbored mutation in this random gene set. We performed 10,000 such random samplings for each pathway and calculated the p-value as the proportion of random samples in which more patients carried mutations than the number of tumors that used the original pathway. The false discovery rate was then calculated for each pathway using the Benjamini and Hochberg method. The significantly enriched pathway was considered if the adjusted p value was less than 0.1.

### Phylogenetic inference

The evolutionary history of each of the 23 tumors were constructed on the basis of the somatic mutations’ reads count using PhyloSub^59^. This approach made use of Bayesian inference and Markov chain Monte Carlo sampling (with 2,500 samplings) to estimate the number of clonal lineages and their ancestry. We only considered trees with the highest likelihood.

### Immunogenic variants prediction

For each somatic point mutation, we obtained the corresponding mutated amino acid and one peptide centered on the mutated residue, flanked on each side by 8 amino acids from the protein sequence. We also obtained the corresponding normal 17-amino-acid peptide. We then used the NETMHC-3.4 algorithm^60^ to predict the binding affinity for the peptide with MHC-I. The variant showed immunogenicity only if the mutated peptide showed strong binding affinity with MHC-I (affinity < 50) and the normal peptide had no binding affinity (affinity > 500) at the same peptide position.

## Acknowledgement

This work was supported by the government funds of Shenzhen, China (SFG 2012.566 and SKC 2012.237), and the National Natural Science Foundation of China (Grant number: 31501065).

## Author contributions

Z-Y.C. developed the concept, designed the research and wrote the manuscript; X. Y. collected the tumor samples; YF. G. and YH. G. performed experiments; X.L. and HL.H. designed the bioinformatics analysis. X.L. performed bioinformatics analysis and drafted the manuscript; HL. H., C-Y. H., and M. Q. contributed to the ideas and edited the paper. All authors read and approved the final manuscript.

## References

1. Brahmer JR, Pardoll DM. Immune checkpoint inhibitors: making immunotherapy a reality for the treatment of lung cancer. Cancer immunology research 1, 85–91 (2013).

2. Borghaei H, et al. Nivolumab versus Docetaxel in Advanced Nonsquamous Non-Small-Cell Lung Cancer. The New England journal of medicine 373, 1627–1639 (2015).

3. Gerlinger M, et al. Intratumor heterogeneity and branched evolution revealed by multiregion sequencing. The New England journal of medicine 366, 883–892 (2012).

4. Nik-Zainal S, et al. The life history of 21 breast cancers. Cell 149, 994–1007 (2012).

5. Waclaw B, et al. A spatial model predicts that dispersal and cell turnover limit intratumour heterogeneity. Nature 525, 261–264 (2015).

6. Chen ZY, Ma F, Huang HL, He CY. Synthetic immunity to breakdown the bottleneck of cancer immunotherapy. Sci Bull 60, 977–985 (2015).

7. Carreira S, et al. Tumor clone dynamics in lethal prostate cancer. Science translational medicine 6, 254ra125 (2014).

8. Eirew P, et al. Dynamics of genomic clones in breast cancer patient xenografts at single-cell resolution. Nature 518, 422–426 (2015).

9. Gundem G, et al. The evolutionary history of lethal metastatic prostate cancer. Nature 520, 353–357 (2015).

10. Couzin-Frankel J. Breakthrough of the year 2013. Cancer immunotherapy. Science 342, 1432–1433 (2013).

11. Nik-Zainal S, et al. Mutational processes molding the genomes of 21 breast cancers. Cell 149, 979–993 (2012).

12. GLOBOCAN 2012: Cervical Cancer Incidence, Mortality and Prevalence Worldwide in 2012. http://globocan.iarc.fr/Pages/factsheetscancer.aspx

13. Ojesina AL, et al. Landscape of genomic alterations in cervical carcinomas. Nature 506, 371–375 (2014).

14. Chung TK, et al. Genomic aberrations in cervical adenocarcinomas in Hong Kong Chinese women. International journal of cancer 137, 776–783 (2015).

15. Cibulskis K, et al. Sensitive detection of somatic point mutations in impure and heterogeneous cancer samples. Nature biotechnology 31, 213–219 (2013).

16. Banerji S, et al. Sequence analysis of mutations and translocations across breast cancer subtypes. Nature 486, 405–409 (2012).

17. Genomes Project C, et al. An integrated map of genetic variation from 1,092 human genomes. Nature 491, 56–65 (2012).

18. Exome Variant Server, NHLBI GO Exome Sequencing Project (ESP), Seattle, WA. http://evs.gs.washington.edu/EVS/. Accessed 4 Jan, 2015

19. Exome Aggregation Consortium (ExAC), Cambridge, MA. http://exac.broadinstitute.org. Accessed 4 Jan, 2015

20. Wright AA, et al. Oncogenic mutations in cervical cancer: genomic differences between adenocarcinomas and squamous cell carcinomas of the cervix. Cancer 119, 3776–3783 (2013).

21. Chung TK, et al. Genomic aberrations in cervical adenocarcinomas in Hong Kong Chinese women. International journal of cancer Journal international du cancer, (2015).

22. Yadav M, et al. Predicting immunogenic tumour mutations by combining mass spectrometry and exome sequencing. Nature 515, 572–576 (2014).

23. Tran E, et al. Cancer immunotherapy based on mutation-specific CD4+ T cells in a patient with epithelial cancer. Science 344, 641–645 (2014).

24. Robbins PF, et al. Mining exomic sequencing data to identify mutated antigens recognized by adoptively transferred tumor-reactive T cells. Nature medicine 19, 747–752 (2013).

25. Trajanoski Z, et al. Somatically mutated tumor antigens in the quest for a more efficacious patient-oriented immunotherapy of cancer. Cancer immunology, immunotherapy : CII 64, 99–104 (2015).

26. Castle JC, et al. Exploiting the mutanome for tumor vaccination. Cancer research 72, 1081–1091 (2012).

27. Guo G, et al. Frequent mutations of genes encoding ubiquitin-mediated proteolysis pathway components in clear cell renal cell carcinoma. Nature genetics 44, 17–19 (2012).

28. Welcker M, Clurman BE. FBW7 ubiquitin ligase: a tumour suppressor at the crossroads of cell division, growth and differentiation. Nature reviews Cancer 8, 83–93 (2008).

29. Mao JH, et al. FBXW7 targets mTOR for degradation and cooperates with PTEN in tumor suppression. Science 321, 1499–1502 (2008).

30. Tommasino M, et al. The role of TP53 in Cervical carcinogenesis. Human mutation 21, 307–312 (2003).

31. Scheffner M, Werness BA, Huibregtse JM, Levine AJ, Howley PM. The E6 oncoprotein encoded by human papillomavirus types 16 and 18 promotes the degradation of p53. Cell 63, 1129–1136 (1990).

32. Scheffner M, Huibregtse JM, Vierstra RD, Howley PM. The HPV-16 E6 and E6-AP complex functions as a ubiquitin-protein ligase in the ubiquitination of p53. Cell75, 495–505 (1993).

33. Boyer SN, Wazer DE, Band V. E7 protein of human papilloma virus-16 induces degradation of retinoblastoma protein through the ubiquitin-proteasome pathway. Cancer research 56, 4620–4624(1996).

34. Jones DL, Thompson DA, Munger K. Destabilization of the RB tumor suppressor protein and stabilization of p53 contribute to HPV type 16 E7-induced apoptosis. Virology 239, 97–107 (1997).

35. McCormack SJ, Brazinski SE, Moore JL, Jr., Werness BA, Goldstein DJ. Activation of the focal adhesion kinase signal transduction pathway in cervical carcinoma cell lines and human genital epithelial cells immortalized with human papillomavirus type 18. Oncogene 15, 265–274(1997).

36. Du M, Fan X, Hong E, Chen JJ. Interaction of oncogenic papillomavirus E6 proteins with fibulin-1. Biochemical and biophysical research communications 296, 962–969 (2002).

37. Chiarugi P, Giannoni E. Anoikis:a necessary death program for anchorage-dependent cells. Biochemical pharmacology 76, 1352–1364 (2008).

38. Nowell PC. The clonal evolution of tumor cell populations. Science 194, 23–28 (1976).

39. Hughes AE, et al. Clonal architecture of secondary acute myeloid leukemia defined by single-cell sequencing. PLoS genetics 10, e1004462 (2014).

40. Hanahan D, Weinberg RA. Hallmarks of cancer: the next generation. Cell 144, 646–674 (2011).

41. Aparicio S, Caldas C. The implications of clonal genome evolution for cancer medicine. The New England journal of medicine 368, 842–851 (2013).

42. Vogelstein B, et al. Cancer genome landscapes. Science 339, 1546–1558 (2013).

43. Moody CA, Laimins LA. Human papillomavirus oncoproteins: pathways to transformation. Nature reviews Cancer 10, 550–560 (2010).

44. Samuels Y, et al. High frequency of mutations of the PIK3CA gene in human cancers. Science 304, 554 (2004).

45. Ciriello G, et al. Emerging landscape of oncogenic signatures across human cancers. Nature genetics 45, 1127–1133 (2013).

46. Rusan M, Li YY, Hammerman PS. Genomic landscape of human papillomavirus-associated cancers. Clinical cancer research 21, 2009–2019 (2015).

47. Dumont AG, Dumont SN, Trent JC. The favorable impact of PIK3CA mutations on survival: an analysis of 2587 patients with breast cancer. Chinese journal of cancer 31, 327–334 (2012).

48. Van Keymeulen A, et al. Reactivation of multipotency by oncogenic PIK3CA induces breast tumour heterogeneity. Nature 525, 119–123 (2015).

49. Liu P, et al. Oncogenic PIK3CA-driven mammary tumors frequently recur via PI3K pathway-dependent and PI3K pathway-independent mechanisms. Nature medicine 17, 1116–1120(2011).

50. Nguyen LV, et al. Barcoding reveals complex clonal dynamics of de novo transformed human mammary cells. Nature 528, 267–271 (2015).

51. Schreiber RD, Old LJ, Smyth MJ. Cancer immunoediting: integrating immunity's roles in cancer suppression and promotion. Science 331, 1565–1570 (2011).

52. Coulie PG, Van den Eynde BJ, van der Bruggen P, Boon T. Tumour antigens recognized by T lymphocytes: at the core of cancer immunotherapy. Nature reviews Cancer 14, 135–146 (2014).

53. Yi X, et al. A new PCR-based mass spectrometry system for high-risk HPV, part I: methods. American journal of clinical pathology 136, 913–919 (2011).

54. Li H, Durbin R. Fast and accurate short read alignment with Burrows-Wheeler transform. Bioinformatics 25, 1754–1760 (2009).

55. McKenna A, et al. The Genome Analysis Toolkit: a MapReduce framework for analyzing next-generation DNA sequencing data. Genome research 20, 1297–1303 (2010).

56. McLaren W, et al. Deriving the consequences of genomic variants with the Ensembl API and SNP Effect Predictor. Bioinformatics 26, 2069–2070 (2010).

57. Van Doorslaer K, et al. The Papillomavirus Episteme: a central resource for papillomavirus sequence data and analysis. Nucleic acids research 41, D571–578 (2013).

58. Subramanian A, et al. Gene set enrichment analysis: a knowledge-based approach for interpreting genome-wide expression profiles. Proceedings of the National Academy of Sciences of the United States of America 102, 15545–15550 (2005).

59. Jiao W, Vembu S, Deshwar AG, Stein L, Morris Q. Inferring clonal evolution of tumors from single nucleotide somatic mutations. BMC bioinformatics 15, 35 (2014).

60. Lundegaard C, et al. NetMHC-3.0: accurate web accessible predictions of human, mouse and monkey MHC class I affinities for peptides of length 8-11. Nucleic acids research 36, W509–512 (2008).

